# Elevated prevalence of the global panzootic chytrid strain in Ecuadorian anurans of the Amazonian lowlands

**DOI:** 10.1101/2024.02.16.580711

**Authors:** Utpal Smart, Shawn F. McCracken, Rebecca M. Brunner, Clarissa Rivera, David Rodriguez

## Abstract

Considerable attention has been directed to studying the infection dynamics of the fungal pathogen, *Batrachochytrium dendrobatidis* (*Bd*), affecting amphibians in the high elevations of the Neotropics. Lowland forests of the same realm, on the other hand, remain relatively understudied in this context. Herein, an attempt to bridge this gap was made by investigating the occurrence of *Bd* in several anuran taxa inhabiting the Amazonian lowlands in the northeast of Ecuador. To this end, 207 anurans belonging to 10 different families, 25 different genera, and 55 distinct host species were sampled for *Bd* DNA in 2008. Data on the taxonomy, morphology (i.e., weight and snout-vent length), and life-long aquatic dependency of hosts (i.e., aquatic index) were also collated to serve as potential predictors of infection prevalence. Genotyping via quantitative PCR revealed the presence of the global panzootic lineage of *Bd* (*Bd-*GPL) in the Ecuadorian Amazon. The overall infection prevalence of *Bd* was determined to be 58%, which is a relatively high prevalence rate of *Bd* reported for any amphibian population from the lowlands of the Neotropics to date. A total of 88% of sampled anuran families tested positive for the infection at varying proportions. A logistic regression analysis showed a significant negative relationship between host weight and the proportion of *Bd* infections (p < 0.05). However, no significant associations were observed between host taxonomy, aquatic dependency, or snout-vent length and *Bd* prevalence. Our findings contribute to the understanding of *Bd* dynamics in the Neotropical lowlands and emphasize the need for future research on the ecological factors influencing *Bd* in the Amazon and their implications for amphibian conservation.

## Introduction

The Neotropics, encompassing Central America, the Caribbean, and South America [1], boast remarkable biodiversity, surpassing the combined diversity of plants and animals in the African and Southeast Asian tropics [2-4]. Unfortunately, this rich biodiversity faces significant risks due to various factors, including global climate change, deforestation, and diseases [5]. Amphibians, particularly anurans, exemplify the precarious state of Central and South American tropical diversity. They constitute one of the most diverse vertebrate groups in the Neotropics [6], yet they are also the most threatened vertebrates, not only in South America but worldwide [7-10]. Among the multitude of threats, emerging infectious diseases (EIDs) have become a major challenge to amphibian conservation in the Neotropics [11,12].

By definition, EIDs are diseases that have either recently appeared within a host population or are rapidly increasing in prevalence or geographic range [1]. EIDs are known to spill over or jump from one taxon to another, thus not just threatening regional biodiversity but whole ecosystems [2-4]. Chytridiomycosis is one such wildlife EID caused by chytrid fungi (i.e., *Batrachochytrium dendrobatidis*, hereafter *Bd*), which infects the skin of amphibians [5-7]. This waterborne disease is characterized by degradation of the mouthparts of larvae or over-keratinization of skin cells in adult amphibians. The resulting failure of gas exchange and electrolyte transport in the animal can eventually lead to death [5,8,9]. *Bd* is known to infect over 700 species [10,11], yet susceptibility to, and prevalence of, this disease seems to be environment, host, population, and strain-specific [12-16].

Amphibians in the Neotropics have been severely impacted by this fungal disease [17,18], and Ecuador is no exception. Ecuador is relevant because it is home to one of the highest numbers of amphibian species in the Neotropics [10,19,20]. Furthermore, over a third of these species are considered amongst the most threatened in South America, as they are being extirpated at an alarming rate due to a host of risk factors which includes *Bd* [20]. Within Ecuador, as in other parts of Central and South America, reports of chytridiomycosis have come from the highlands [21-23]. Meanwhile, the relevance of lowland habitats, such as the Amazon Basin, often tends to be underrepresented in studies on the ecology and epidemiology of *Bd* [24,25]. Thus, major knowledge gaps still exist about strain-specific disease dynamics and their implications in these low-elevation regions of South America [25,26].

Genotyping of *Bd* from Ecuador is still needed, despite confirmation of infected individuals from the highlands of the Andes [20,22] and the lowlands of the Amazon [27]. This is important given that there are multiple divergent lineages of *Bd* currently recognized. Three distinct strains have been detected in South America, namely, the panzootic lineage called *Bd-*GPL, the *Bd-*Asia-2/*Bd-*Brazil lineage, and a hybrid strain. *Bd-*GPL, the most prevalent and hypervirulent among all the strains [14,28], has been detected in at least five different South American countries; specifically, Colombia, Peru, Brazil, Chile, and Bolivia [29]. The *Bd*-Asia-2/*Bd*-Brazil lineage has been reported in the Atlantic Forest of Brazil and even detected on a frog from a local market in Michigan, United States, which originated from Brazil [39]. The diversity and distribution of these strains underscore the need for further genotyping studies in Ecuador to better understand the prevalence and dynamics of *Bd* infections.

In addition to genotyping *Bd*, understanding the influence of host traits (e.g., behavior, size, and life history) on disease susceptibility in the Ecuadorian Amazon also needs more attention. For instance, in parts of South and Central America, the effects of *Bd* are found to be correlated to the host’s exposure to water during different life-history stages [13,15,30,31]. Accordingly, direct-developing (i.e., metamorphosis absent) species that have little or no contact with aquatic environments during their ontogeny [32,33], generally have lower *Bd* prevalence than their aquatic counterparts [13,15,34,35] (see [36,37] for exceptions). It is to be noted, however, that most of these data come from studies focusing on the highlands. Whether the same underlying factors apply to host-pathogen relationships in the lowlands of the Amazon Basin still requires investigation.

Our study, aimed to (1) genotype *Bd* collected from various anuran species found in a lowland Amazonian Forest in Ecuador; and (2) investigate the association between aquatic dependency, morphology, familial taxonomy, and prevalence of *Bd.* In doing so, we expand ongoing attempts at divulging crucial information on the obscure infection patterns of the amphibian-killing fungus in Neotropical lowlands.

Our results reveal a high prevalence of *Bd-*GPL among the Amazonian lowland anuran fauna of Ecuador. Additionally, our study highlights the need to reconsider previously identified predictors of *Bd* dynamics, such as aquatic dependency, size, and taxonomy, in the context of lowland tropical forests, with special attention to bromeligenous species. Although major *Bd*-driven amphibian declines have not been reported in lowland South American species, our results suggest that these sites could still contribute to the spread and persistence of chytrid.

## Materials and methods

### Study Site

Samples were originally collected at the Tiputini Biodiversity Station, Orellana Province, Ecuador (-0.637859°S, -76.149823°W, 217 m Elev.). Founded in 1994 by Universidad San Francisco de Quito (USFQ), the station lies along the Rio Tiputini and is adjacent to the Yasuni Biosphere Reserve, which is renowned for its biodiversity [38]. The research station has 139 documented amphibian species within its 6.5 km^2^ boundary, spanning the three orders Caudata, Gymnophia, and Anura, with 150 species known from the greater Yasuni Biosphere Reserve [38,39].

### Sample collection

All observations and data were collected between May-Aug 2004, May-August 2006, and April-Nov 2008. We used archived toe and thigh skin clips from anurans originally collected near the forest floor to the upper canopy. DNA was extracted using the DNeasy Blood & Tissue Kit (Qiagen, Inc.), and DNA presence and quality were assessed using agarose electrophoresis. Morphological measurements for snout-vent length (SVL) were taken with a Mitutoyo CD-S6”C digital dial caliper (0.01 mm precision) and weights were taken with a Pesola digital pocket scale (0.01 g precision) on live specimens. Research was conducted in compliance to the rules overseen by the Texas State University Institutional Animal Care and Use Committee (Permit #: 0721-0530-7, 05-05C38ADFDB, and 06-01C694AF). Permission and permits issued by the Ministerio del Ambiente, Ecuador (Permit #: 006-IC-FA-PNY-RSO, 012-IC-FA-PNY-RSO, Provincial de Orellana Fauna permit number 0018 DPO-MA, and Provincial de Napo Fauna permit number 017-IC-FAU/FLO-DPN/MA).

### Quantitative polymerase chain reaction (qPCR) for Bd genotyping

Reactions for the quantification of *Bd* load were run in singlicate 25 µL volumes comprising 5 µL of DNA (diluted 1:10) and 12.5 µL of TaqMan Fast Advanced Master Mix (Thermo Fisher Scientific, Inc.), 2.75 µL nuclease-free H_2_O, 0.625 µL of primer ITS1-3 Chytr (18 µM), 0.625 µL of primer 5.8S Chytr (18 µM), 0.625 µL of probe Chytr MGB2 (5 µM), and 0.50 µL bovine serum albumin (400 ng/µL) per reaction [40]. A standard curve was generated using the global panzootic strain JEL423 [41], which had a dynamic range of 0.1 to 1,000 zoospore equivalents (ZE). We considered samples *Bd* positive via qPCR if the load was greater than 1 ZE.

All samples were genotyped, regardless of *Bd* presence or absence, using a single nucleotide polymorphism (SNP) assay that discriminates between *Bd*-GPL and *Bd*-Asia2/*Bd*-Brazil (SC9_200709_CT) based on 27 global genomes [42]. The primers amplify a 109 base pair fragment, and dual probes target a SNP at position 200,709 on the supercont1.9 genomic scaffold of the strain JEL423 reference genome (GenBank: DS022308.1). The dual probes can detect either *Bd*-GPL (genotype TT), *Bd*-Asia2/*Bd*-Brazil (genotype CC), or a co-infection or hybrid strain (genotype CT) (Table 1). Genotyping reactions were conducted in singlicate 15 µl volumes comprising of 15 µl of TaqMan Fast Advanced Master Mix, 0.75 µl of the SNP assay (20X concentration), 4.25 µl of nuclease-free H2O, and 5 µl of extracted DNA (variable concentrations). The results were interpreted using the Thermo Fisher Connect cloud service and the “Standard Curve” and “Genotyping” applications to detect *Bd* presence/absence, *Bd* infection load, and generate the *Bd* SNP genotype calls.

**Table 1.**
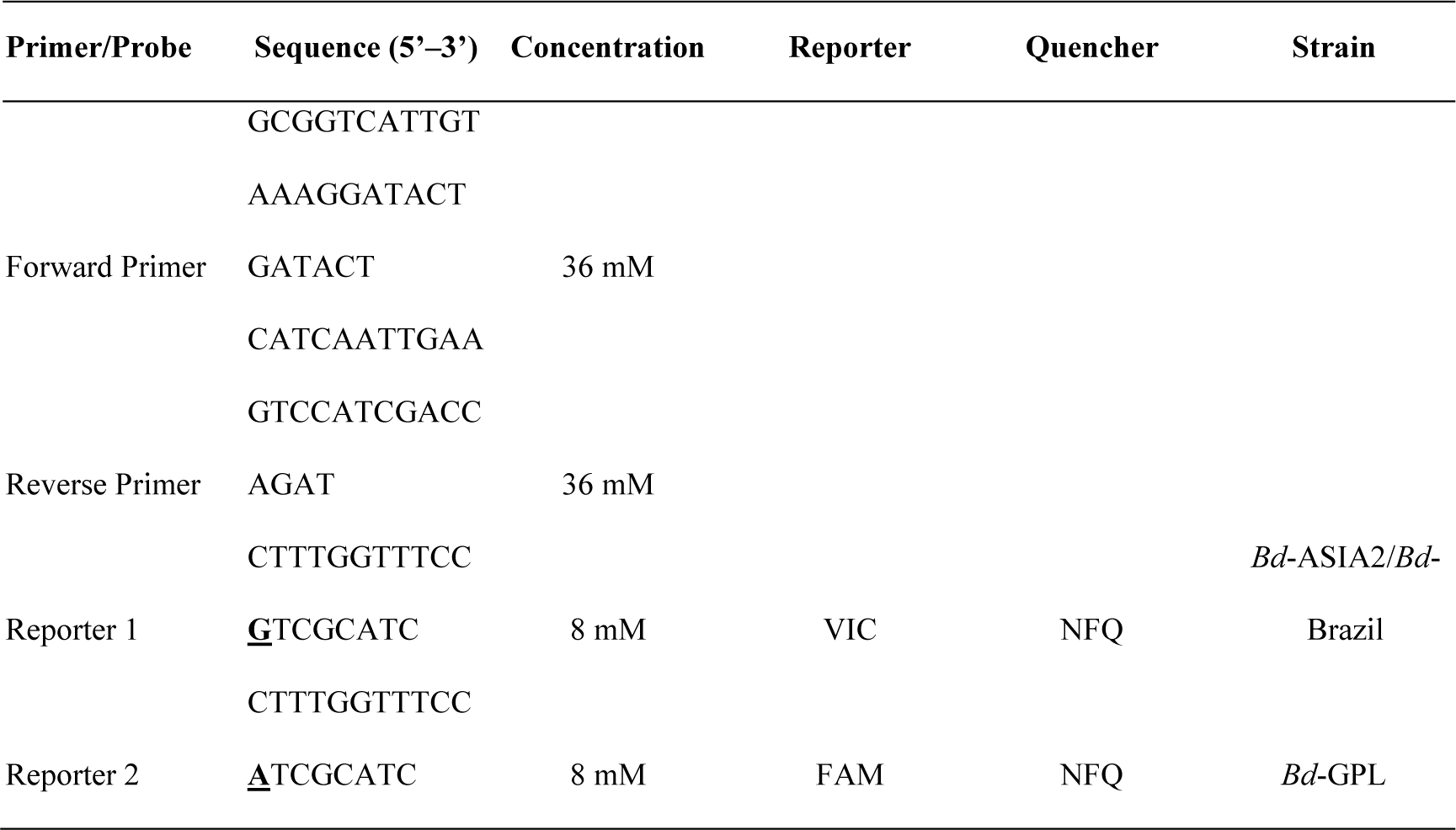
Primer and probe sequences for SC9_200709_CT (Assay ID AHGJ91E), a custom TaqMan SNP genotyping assay (Applied Biosystems, Inc.) at 40X concentration (SNP in bold and underlined). This assay targets the nuclear genome of *Batrachochytrium dendrobatidis* s and discriminates between alleles that identify strains *Bd*-GPL or *Bd*-ASIA2/*Bd*-Brazil.

### Statistical analyses

Before conducting the statistical analyses, rows with missing data (i.e., NA) were excluded from the dataset to ensure the accuracy of the computations. All analyses were conducted in the R environment for statistical computing (version 4.3.0) [43].

The overall prevalence of *Bd* was determined by calculating the ratio of samples that tested positive for *Bd* to the total sample size. Additionally, the prevalence was calculated for each family of anuran sampled, and 95% Wilson binomial confidence intervals (CI) were generated for prevalence estimates using the epi.conf() function in the epiR package [54]. To maximize overall sample size and account for the uncertain species-level identification of some samples, individuals were grouped by taxonomic family. Summary plots were created using the package ggplot2 [55] for clear visualization of the results.

To investigate the relationship between infections and host traits, we performed a binomial logistic regression analysis. Specifically, we aimed to test the null hypothesis that there is no significant relationship between the taxonomy (at the family level), morphology (SVL and weight), aquatic index (AI) of the host, and the probability of *Bd* infections. AI assignments were based on the degree of exposure to water during different life history stages of each sampled anuran family, following the approach of [56] with some modifications, references to literature sources such as AmphibiaWeb [43,57], and the IUCN Red List of Threatened Species [10]. Anuran families were categorized into four AI categories: AI0 for terrestrial species with direct development (terrestrial breeders), AI1 for arboreal species that breed in water, AI2 for riparian species that breed in water, and AI3 for direct-developing bromeligenous species, a new category introduced in this study to encompass species that rely heavily on the moist microhabitat of phytotelmata for shelter and/or breeding, bypassing the aquatic larval stage (pers. obs.) [25,35,40].

Before conducting the logistic regression, we examined potential extreme outliers in the dataset using Cook’s distance via the cooks.distance() function. After confirming the absence of extreme outliers, we ran the initial regression model using the base R function glm() with all four explanatory variables. The results of this regression were then subjected to the vif() function implemented in the car package [44] to check for multicollinearity using generalized variance inflation factors (GVIF). We observed significant multicollinearity between the Family and AI variables, as well as weak multicollinearity between SVL and Weight. To address this issue, we removed the Family variable from the regression model and log-transformed the two continuous morphological variables to mitigate the effects of multicollinearity. Subsequently, we conducted a final logistic regression with three explanatory variables. We plotted the odds ratio plot for this regression analysis using the function or_plot() in the package finalfit [45].

## Results

A total of 207 individual anurans were sampled, spanning 9 families, 25 genera, and approximately 55 known species of anurans (Supplementary Table S1). All DNA extractions from the toe clips showed a high molecular weight band on an agarose gel. Out of 120 *Bd*-positive samples via qPCR, 72 (60%) were genotyped as strain *Bd*-GPL based on the results of the SNP Assay. The remaining 48 positive samples did not show an amplification curve for either dye. As expected, samples that were qPCR negative also did not return a genotype. The median infection intensity for genotyped samples was 2,329 ZE with a range of 458–1,048,416 ZE. The median infection intensity for non-genotyped samples was 122 ZE with a range of 1.83–2,656 ZE (Supplementary Figure S1). The difference between the two median infection intensity values was significantly different (two-sample Wilcoxon test; W = 306, α = 0.05, p-value = 2.2e-16).

The overall prevalence of *Bd* infections was 58.0 % (95% CI = 0.51–0.64; n = 207). When grouped by taxonomic family, the single representatives of *Aromobatidae* and *Ranidae* tested positive, while the single representative of *Centrolenidae* tested negative for *Bd* (Table 2, Figure 1). When grouped by AI, frogs belonging to AI2 had the highest prevalence of chytrid at 64% (95% CI = 0.49–0.76; n = 44), followed by a prevalence of 57% for groups AI0 (95% CI = 0.48–0.67; n = 65) and AI3 (95% CI = 0.41-0.72; n = 35), and AI1 with an infection prevalence of 54% (95% CI = 0.42–0.66; n = 63) (Table 3).

**Figure 1.**
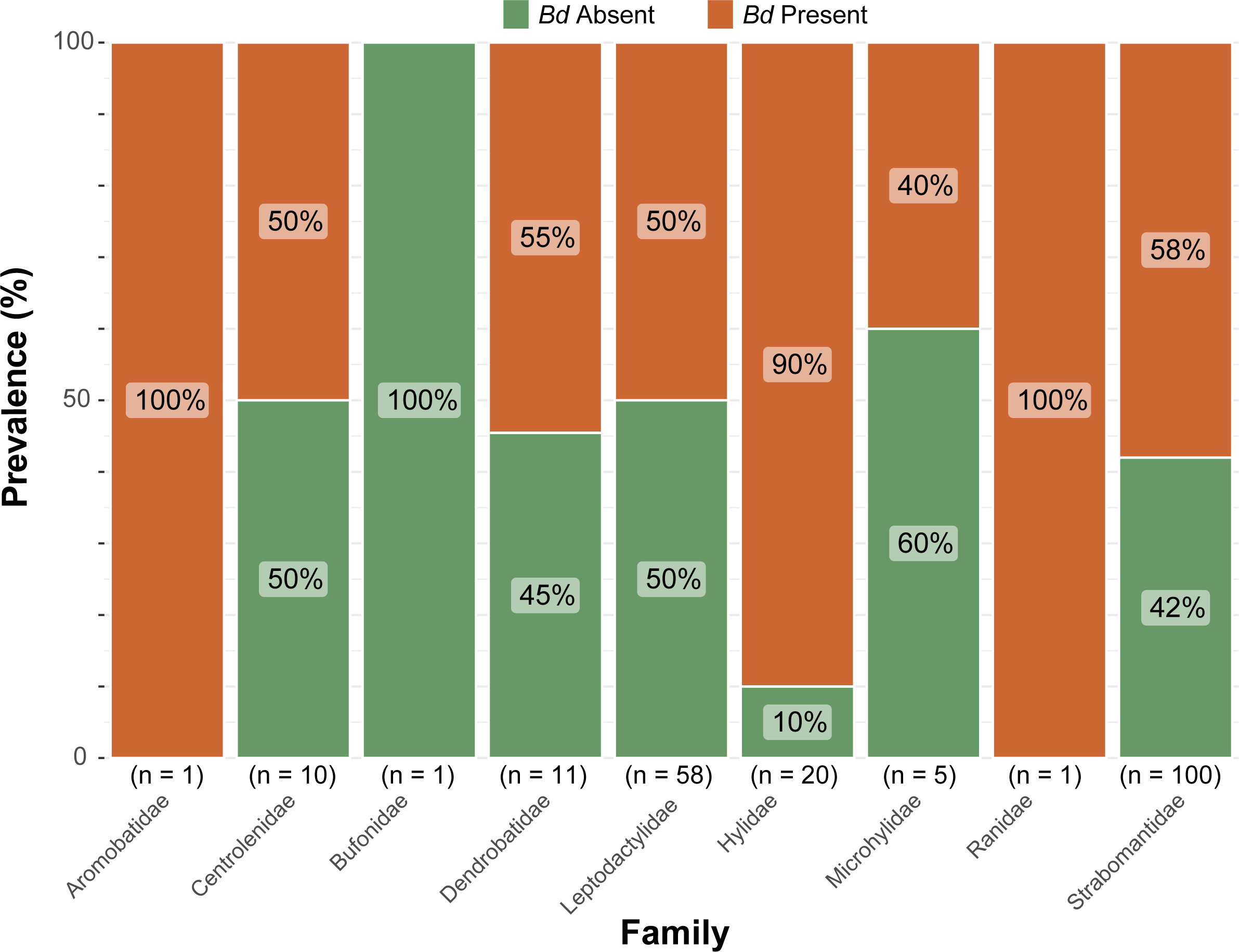
Prevalence of *Batrachochytrium dendrobatidis* (*Bd*) infections in nine different families of anurans found at Tiputini Biodiversity Station. The x-axis displays the frog families and their respective sample sizes (n), while the y-axis represents the *Bd* prevalence in percentages (%). The colours indicate *Bd*’s presence (reddish orange) and absence (olive green). Refer to Table 2 for confidence intervals of the prevalence.

**Table 2.**
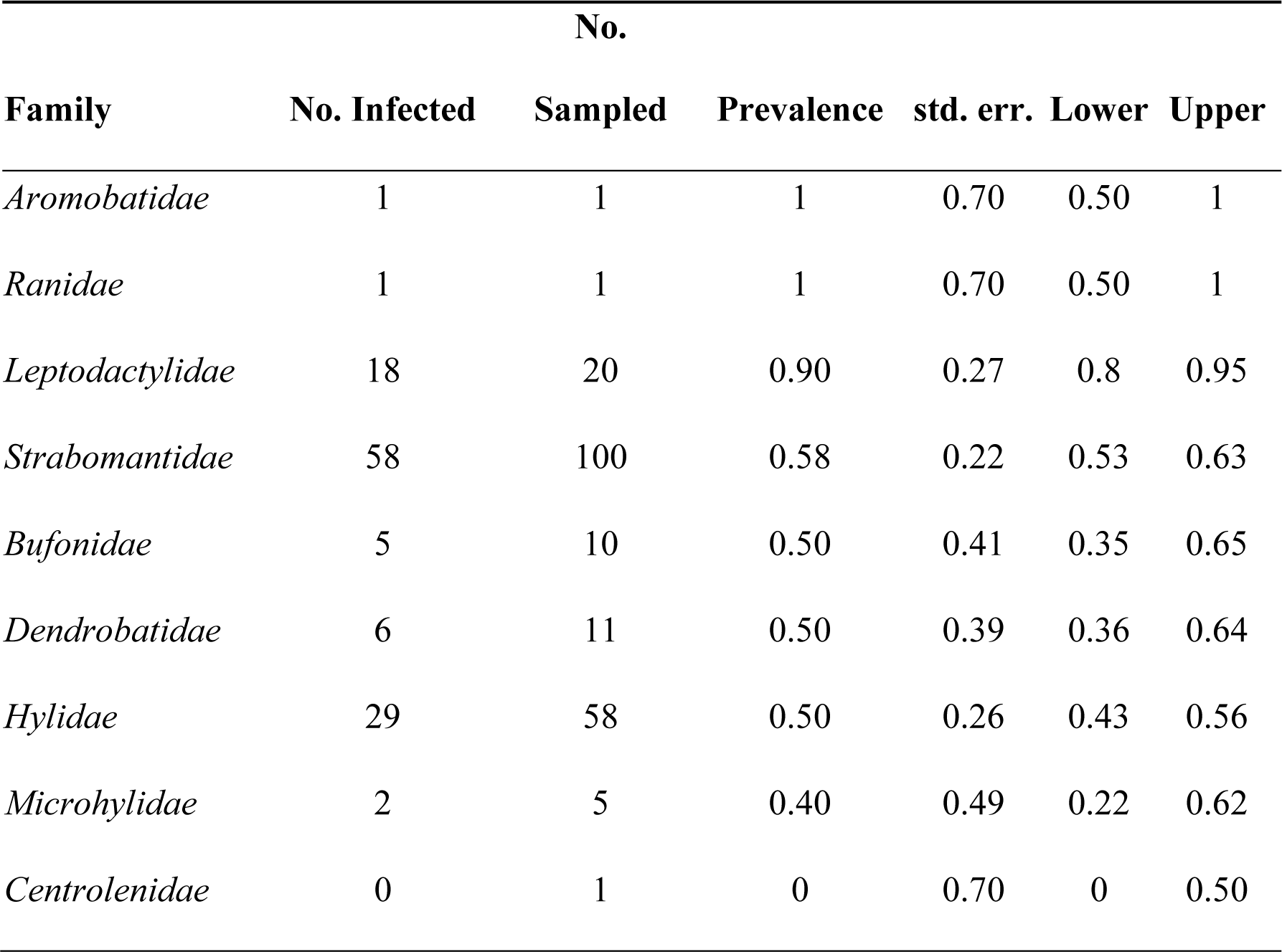
Prevalence of *Batrachochytrium dendrobatidis* (*Bd*) infections in Tiputini Biodiversity Station sorted in descending order and partitioned into the nine taxonomic families that were sampled. Columns represent the number of infected individuals (“No. Infected”), the total number of individuals sampled (“No. Sampled”), mean prevalence (“Prevalence”), standard error ("std. err.”), and lower and upper 95% Wilson binomial confidence intervals (“Lower” and “Upper” respectively).

**Table 3.** Prevalence of *Batrachochytrium dendrobatidis* (*Bd*) on the anurans of Tiputini Biodiversity Station sorted in descending order and partitioned by four aquatic index (AI) categories. AI0 represents terrestrial species with terrestrial eggs, AI1 represents arboreal species with aquatic larvae, AI2 represents terrestrial species with aquatic larvae, and AI3 represents arboreal species with terrestrial eggs (or non-aquatic larvae). Columns represent the number of infected individuals (“No. Infected”), the total number of individuals sampled (“No. Sampled”), mean prevalence (“Prevalence”), standard error (“std. err.”), and lower and upper 95% Wilson binomial confidence intervals (“Lower” and “Upper” respectively).

| AI | No. Infected | No. Sampled | Prevalence | std. err. | Lower | Upper |
| --- | --- | --- | --- | --- | --- | --- |
| 0 | 38 | 65 | 0.57 | 0.25 | 0.46 | 0.70 |
| 1 | 34 | 63 | 0.54 | 0.25 | 0.42 | 0.66 |
| 2 | 28 | 44 | 0.64 | 0.27 | 0.49 | 0.76 |
| 3 | 20 | 35 | 0.57 | 0.30 | 0.41 | 0.72 |

The results of the logistic regression revealed that weight had a significant negative effect on the probability of *Bd* infections (β = -0.42, p = 0.02*) (Table 4, Figure 3). However, SVL (β = 0.02 p = 0.96) and the AI categories (AI0 [intercept] β = 0.25, p = 0.85; AI1: β = 0.41, p = 0.31; AI2: β = 0.55, p = 0.22; AI3: β = 0.03, p = 0.94) did not show significant effects on the likelihood of *Bd* infections (Table 4, Figure 3).

**Table 4.** Results of the logistic regression investigating the relationship between host traits (“Term”) and *Batrachochytrium dendrobatidis* (*Bd*) infections in anurans from Tiputini Biodiversity Station. The “Estimate” column provides the coefficient estimates for each predictor in the logistic regression model. The “std. err.” column shows the standard errors associated with each coefficient estimate. The “Statistic” column displays the z-value (Wald statistic) for each predictor. The "p.value" column presents the p-values associated with each coefficient estimate. Note that the term “Weight” was found to be significant (p < 0.05) and is highlighted by an asterisk(*).

| Term | Estimate | std. err. | Statistic | p.value |
| --- | --- | --- | --- | --- |
| (Intercept) | 0.25 | 1.32 | 0.19 | 0.85 |
| AI1 | 0.41 | 0.40 | 1.02 | 0.31 |
| AI2 | 0.55 | 0.45 | 1.21 | 0.22 |
| AI3 | 0.03 | 0.44 | 0.08 | 0.94 |
| Weight* | -0.42 | 0.19 | -2.27 | 0.02 |
| SVL | 0.02 | 0.41 | 0.04 | 0.97 |

## Discussion

Since chytrid infections in amphibians first gained widespread attention in the 1990s [11,46,47], a significant amount of research has been carried out on various aspects of chytridiomycosis in the Neotropics. However, the majority of these *Bd* studies, thus far, have prioritized highlands and montane ecosystems over the biodiverse lowland tropical forests. While these warm lowlands might not provide abiotic conditions within the optimal physiological parameters for this pathogenic fungus [23,24,48,49] (however, see McCracken et al [27] for exception), the role of Neotropical lowlands as putative *Bd* reservoirs or sinks has only recently begun to be investigated [25,50-52]. Our study is one amongst only a handful of contributions to a better understanding of the associations between anuran host traits and chytrid infections in the lowlands of Central and South America.

### Genotyping and overall Bd prevalence

To the best of our knowledge, no prior information on the genotype of *Bd* was known from this region of Ecuador. Past published assessments provided only presence/absence data for *Bd* in the Ecuadorian Amazon [22,27]. Based on our results, this site only showed evidence of the global panzootic lineage with no presence of *Bd* - Brazil/Asia2. This is consistent with the genotyping results for *Bd* from the Peruvian Amazon [51] and extends the range of *Bd*-GPL to now include Ecuador along with other South American countries like Brazil, Chile, and Colombia.

In the context of the overall prevalence of chytrid infections, our results indicate a higher prevalence (58.0%; n = 207) than those reported by earlier studies from the Amazonian lowlands (3.8%; n = 1391 [25], 34.0%; n = 324 [51], and 0.7-7.3%; n = 282 [53]), including one that was carried out at the same location by McCracken et al. [27] (20.0%; n = 86). However, our prevalence data are comparable to other studies in the lowlands of Costa Rica in Central America (54.6%; n = 348 [50]) and Brazil in South America [54]. Notably, our study records the highest overall prevalence of chytrid recorded from Amazonian lowlands to date, showing that lowlands of the Ecuadorian Amazon, which have been historically understudied in *Bd* research, warrant closer attention.

On a more cautionary note, our molecular sampling results show a clear and strong positive association between the infection intensity (i.e., the load of *Bd*) and odds of the chytrid strain being successfully genotyped (Supplementary Figure S1). This implies that samples that tested positive for *Bd* might not have returned a genotype unless the fungal load was above a given threshold (roughly between 458 ZE and 2,656 ZE; Supplementary Figure S1). Future studies should be mindful that *Bd* infection prevalence rates in a sampled population are prone to underestimation if single SNP qPCR genotyping is used as the sole method of chytrid detection.

### Body measurements and Bd prevalence

According to our results, the weight of the host was found to be significant in relation to the presence/absence of *Bd.* The median weight of uninfected individuals (1.80 g) was slightly larger than the median weight of infected individuals (1.50 g) (Table 4, Figures 2a and 3). This result suggests that for each unit increase in weight, the log-odds of being infected with *Bd* decrease by 0.42207, indicating that larger individuals might be less susceptible to *Bd* infections compared to their smaller counterparts. On the other hand, the median SVL of uninfected anurans (27.60 mm) was also higher than that of infected anurans (26.95 mm) (Figure 2b) but did not show a statistically significant effect on *Bd* infections (p-value = 0.9656) (Table 4, Figures 2b and 3).

**Figure 2.**
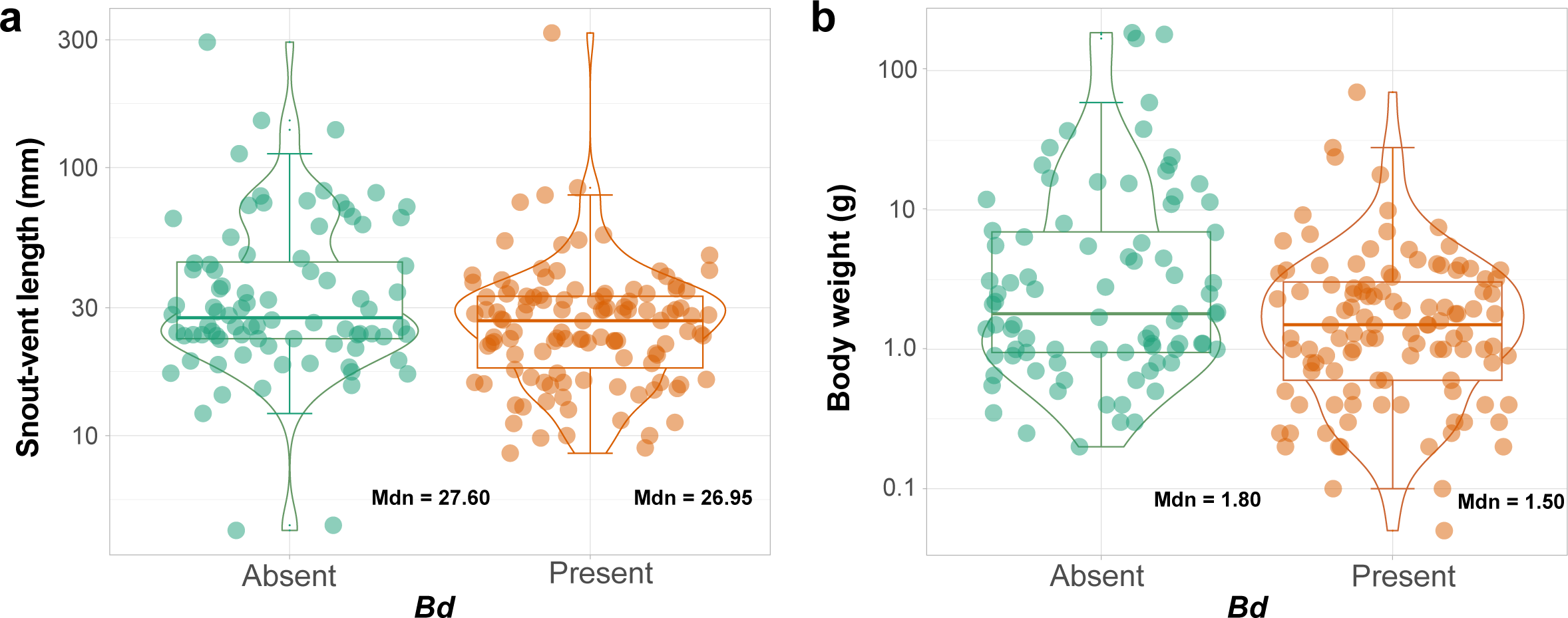
Combined dot plots, boxplots, and violin plots comparing snout-vent length (SVL) 2a and weight 2b between frogs infected with *Batrachochytrium dendrobatidis* (*Bd* present) and non-infected (*Bd* absent) frogs in the dataset. The x-axis represents the state of infection and the (log-transformed) y-axis represents SVL in millimeters (mm) on the left and weight in grams (g) on the right. The box represents the interquartile interval. The vertical line in the middle of the box represents the median (Mdn). The whiskers represent the maximum and minimum values of the body measurements.

**Figure 3.**
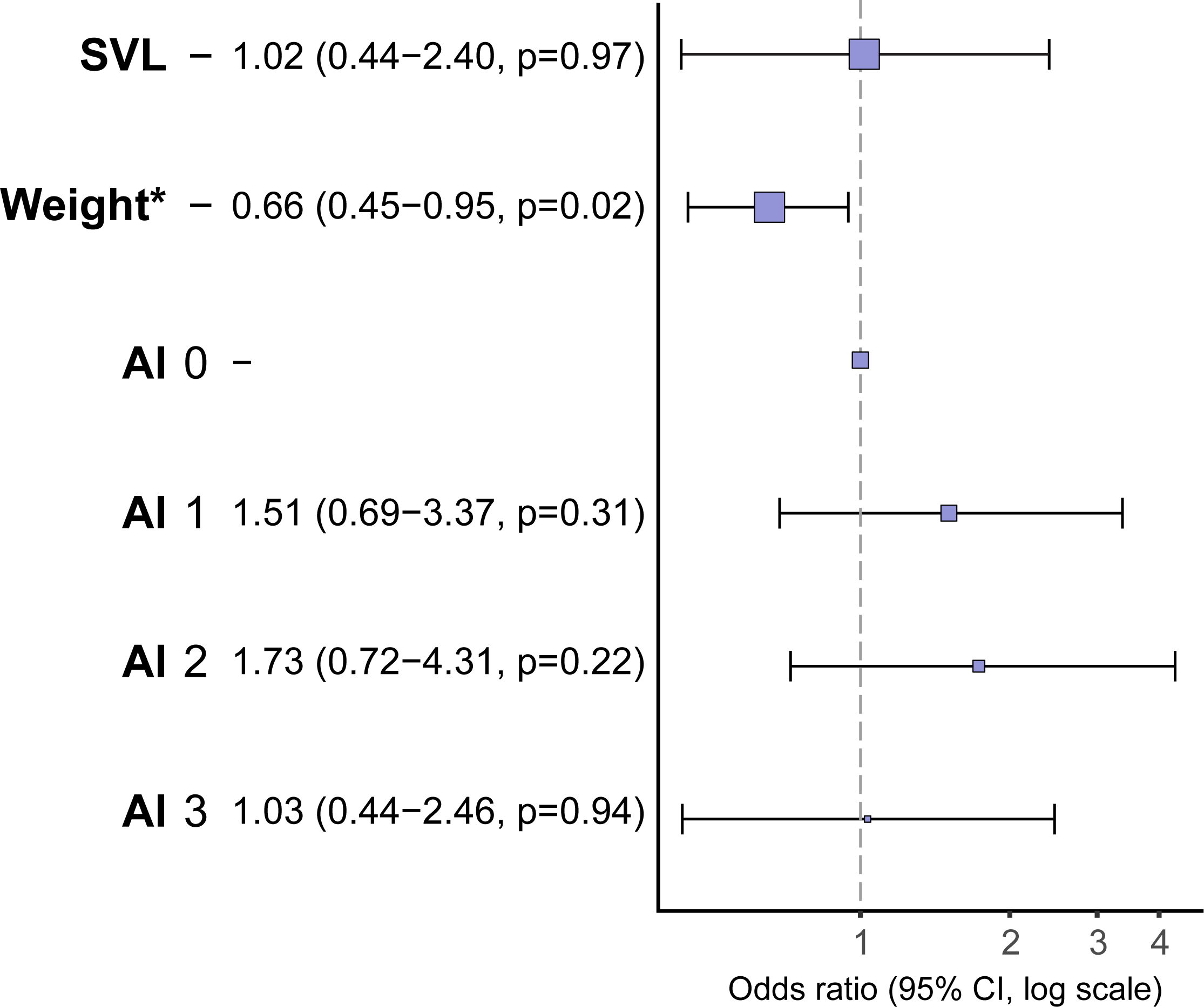
Odds Ratio Plot for the Logistic Regression Results, illustrating the significance of predictors on *Batrachochytrium dendrobatidis* (*Bd*) infections in anuran species at Tiputini Biodiversity Station. The plot displays the odds ratios, along with their corresponding confidence intervals and p-values, for the variables "Weight," "Snout-Vent Length (SVL)," and the four categories of "Aquatic Index (AI)." The predictor "Weight" is identified as significant as indicated by the asterisk (*).

The available literature is conflicted on the influence of host size on the infection intensity and prevalence of *Bd*. Some research has shown that larger (and older) hosts in several different organisms (including frogs infected with *Bd*) can have more developed immune systems and are therefore able to mount better defenses against pathogens [55-57]. For example, Burrow et al. [58] investigated the association between host size and *Bd* and found that smaller size in anurans increased susceptibility to diseases. Similarly, studies on Australian frogs have uncovered an inverse relationship between the likelihood of chytrid infection and SVL [15,40]. Conversely, research has also shown that larger hosts are not only more likely to be infected but also more likely to experience decline [12,13]. This correlation between an anuran’s size/weight and its ability to combat *Bd* infections appears to be more complex than a simple linear relationship between the two variables. For example, according to Lips et al. [13], large frogs infected by *Bd* only declined in high elevations, whereas large infected lowland anurans survived. This finding hints at the existence of some form of interaction between traditional predictors of *Bd* in their effect on the prevalence and intensity of the pathogen.

The relationship between body measurement and *Bd* infections in our data underscore the need to investigate whether size and weight drive the anuran’s abilities to prevent infections, or conversely, whether infections pose constraints on how large an anuran can grow. There is support for both hypotheses in the literature [56,59], and clearly, more data are required before these questions can be addressed. For now, our results show a more pronounced prevalence of *Bd* infections in smaller anuran species, from an important ecological site within the Ecuadorian Amazon, compared to larger taxa.

### Aquatic indices, taxonomy, and Bd prevalence

Our logistic regression analysis revealed no significant association between the AI and the prevalence of chytrid infections in anurans from Tiputini (Table 3). Anurans with an AI2 had the highest *Bd* prevalence (64%), while those with an AI1 had the lowest prevalence (54%), slightly lower than individuals with an AI0 and AI3 (57%).

Previous studies [12,13,15,30] have found aquatic breeders (AI2) to have a statistically higher prevalence of *Bd* especially when compared to terrestrial breeders. It is worth noting, however, that this does not appear to be a consistent pattern across all studies. For example, Ribeiro et al. [36], recorded a higher *Bd* prevalence for direct-developing/terrestrial frogs compared to aquatic breeders. The authors of that study accredited this to the fact that, unlike other research, their sampling was restricted to riparian zones, which are areas that may facilitate contact with waterborne chytrids regardless of the type of breeding environment [35,36,60]. Stream-adjacent populations of direct-developing frogs could thus be at a higher risk of infection by *Bd* than currently thought. This reinforces the need for more surveys that focus on these lowland riparian environments and their role in the host-pathogen dynamics of chytrid.

Though they did not investigate the association between exposure to water and *Bd* infections, McCracken et al. [27] did find that prevalence of chytrid was non-randomly distributed along the vertical axis, i.e., the frogs that inhabited the canopy (defined as over 4 meters above ground level) had the highest prevalence of *Bd* with as many as 33% being infected. This further corroborates our findings of lack of association between AI and *Bd* prevalence, considering that several canopy inhabiting species in our dataset are either direct developers or terrestrial species that lay eggs in water (i.e., AI0 and AI2, respectively).

It has been proposed that canopy-dwelling species may be exposed to chytrid fungi present in stagnant water collected within the phytotelmata of tank bromeliads phytotelmata of tank bromeliads [27]. Several species of frogs are known to exploit these water-filled plant cavities for egg or tadpole development or to even spend their entire life cycle within them [61-63]. To factor in the near-constant exposure to water or humidity, species occupying this niche (all belonging to the genus *Pristimantis*) sampled in our study, were assigned a separate AI category (AI3) than non-bromeligenous direct developers. Indeed, the prevalence of *Bd* was higher in these bromeligenous frogs when compared to aquatic breeders that were arboreal. This is not entirely surprising, since even though ambient temperatures in lowland tropical rainforests may not be optimal for the proliferation of *Bd*, McCracken et al. [27] found the water in lowland bromeliads to be at temperatures that are conducive for the survival of *Bd*. A high prevalence of *Bd* has also been reported for frogs inhabiting phytotelma microhabitats in other Neotropical lowland forests [64]. Future chytrid research would thus benefit by investigating the role of this water-impounding foliage as reservoirs of *Bd* in lowland tropical forests.

Given the high correlation observed between taxonomic rank at the family level and aquatic indices in our dataset, we are confident in extrapolating that the prevalence of the disease was likely randomly distributed among taxonomic families. While contradictory to some previous findings [12,65], this lack of association is documented by others that have looked at phylogenetic relatedness as a predictor of *Bd* prevalence and susceptibility [11,66]. Two families had an infection prevalence of 100%, namely *Ranidae* (n=1) and *Aromobatidae*, (n=1); however, greater sample sizes are certainly needed for these taxa. *Strabomantidae* (n=100), the family of direct developers with no association with water for breeding (i.e., AI0 and AI3) in the dataset, had a *Bd* prevalence of 58% albeit with higher accuracy (95%, CI 0.53-0.63) given the larger sample size (Table 2).

*Centrolenidae* (n = 1) had no *Bd* prevalence with the caveat that only one individual was detected and that this family also requires additional sampling at this site (Table 2). The lowest non-zero mean of *Bd* prevalence was as high as 40% (95% CI 0.22-0.62) in the *Microhylidae* (n = 5), yet more sampling is also needed for this group. All remaining prevalence estimates were at or above 50%, with *Leptodactylidae* (n = 20) showing a 90% (95% CI 0.80-0.95) prevalence of *Bd* (Table 2). Overall, 88% (8 out of 9) of the families sampled for *Bd* were found to be infected by the fungus, compared to 43% (3 out of 7) of the amphibian families that were sampled by McCracken et al. [27] at the same site. Similar to our findings regarding morphology, our results do not align with existing literature from studies conducted in the highlands of the Neotropics concerning the association between taxonomy and the prevalence of *Bd* infections in anuran fauna.

## Conclusion

Our study contributes to the limited body of research investigating the prevalence of *Bd* infections in the lowland Amazonian rainforest of Ecuador [11,33,37]. Notably, we confirm the presence of the *Bd*-GPL strain infecting amphibians in this region and report a comparatively high prevalence of *Bd* infections among Neotropical lowland anuran fauna. Furthermore, our findings highlight the need to re-evaluate previously identified predictors of *Bd* dynamics, such as morphology, aquatic dependence, and taxonomy, which were primarily derived from studies focused on highland environments. In the context of lowland tropical forests, special attention should be given to bromeligenous species. Although major *Bd*-driven amphibian declines have not been reported in lowland South American species, our results suggest that these sites could still contribute to the spread and persistence of chytrid, a pathogen responsible for one of the most devastating wildlife EIDs in modern history.

## FIGURE CAPTIONS

**Figure S1.**
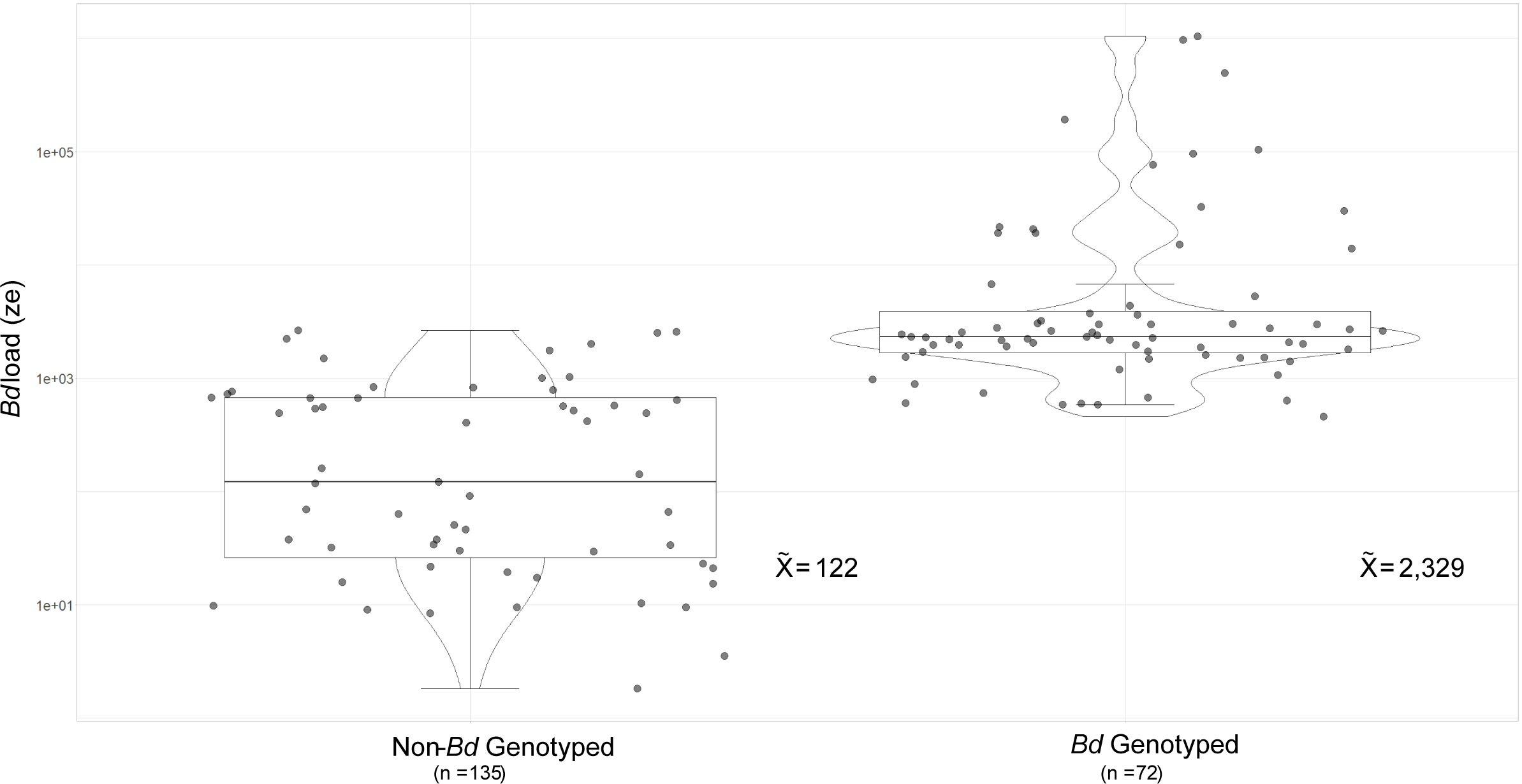
Combined dot plots, boxplots, and violin plots comparing the distribution of *Bd* load in non-genotyped vs. non-genotyped frogs in the dataset. The x-axis represents the genotyping status. The (log-transformed) y-axis represents the *Bd* load in zoospore equivalents (ZE). The box represents the interquartile interval. The vertical line in the middle of the box represents the median 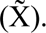 The whiskers represent the maximum and minimum values of the load.

## References

1. World Health Organization. Regional Office for South-East A. A brief guide to emerging infectious diseases and zoonoses. New Delhi: WHO Regional Office for South-East Asia; 2014. en.

2. Hoberg EP, Brooks DR. Evolution in action: climate change, biodiversity dynamics and emerging infectious disease. Phil Trans R Soc B. 2015;370(1665):20130553.

3. Borremans B, Faust C, Manlove KR, et al. Cross-species pathogen spillover across ecosystem boundaries: mechanisms and theory. Phil Trans R Soc B. 2019;374(1782):20180344.

4. Daszak P, Cunningham AA, Hyatt AD. Emerging infectious diseases of wildlife--threats to biodiversity and human health. Science. 2000;287(5452).

5. Grogan LF, Robert J, Berger L, et al. Review of the amphibian immune response to chytridiomycosis, and future directions. Front Immunol. 2018;9.

6. Martel A, Pasmans F, Fisher MC, et al. Chytridiomycosis. In: Seyedmousavi S, de Hoog GS, Guillot J, et al., editors. Emerging and Epizootic Fungal Infections in Animals. Cham: Springer International Publishing; 2018. p. 309-335.

7. Scheele BC, Pasmans F, Skerratt LF, et al. Amphibian fungal panzootic causes catastrophic and ongoing loss of biodiversity. Science. 2019;363(6434).

8. Skerratt LF, Berger L, Speare R, et al. Spread of chytridiomycosis has caused the rapid global decline and extinction of frogs. EcoHealth. 2007;4(2):125.

9. Van Rooij P, Martel A, Haesebrouck F, et al. Amphibian chytridiomycosis: a review with focus on fungus-host interactions. Vet Res. 2015;46(1):137.

10. Frost DR, Grant T, Faivovich J, et al. The amphibian tree of life. Bull Am Mus Nat Hist. 2006;2006(297):1–291, 291.

11. Berger L, Speare R, Daszak P, et al. Chytridiomycosis causes amphibian mortality associated with population declines in the rain forests of Australia and Central America. Proc Natl Acad Sci U S A. 1998;95(15):9031–9036.

12. Bancroft BA, Han BA, Searle CL, et al. Species-level correlates of susceptibility to the pathogenic amphibian fungus *Batrachochytrium dendrobatidis* in the United States. Biodivers Conserv. 2011;20(9):1911–1920.

13. Lips KR, Reeve JD, Witters LR. Ecological traits predicting amphibian population declines in central america. Conserv Biol. 2003;17(4):1078–1088.

14. James TY, Toledo LF, Rödder D, et al. Disentangling host, pathogen, and environmental determinants of a recently emerged wildlife disease: lessons from the first 15 years of amphibian chytridiomycosis research. Ecol Evol. 2015;5(18):4079–4097.

15. Kriger KM, Hero J-M. The chytrid fungus *batrachochytrium dendrobatidis* is non-randomly distributed across amphibian breeding habitats. Divers Distrib. 2007;13(6):781–788.

16. Savage AE, Mulder KP, Torres T, et al. Lost but not forgotten: MHC genotypes predict overwinter survival despite depauperate MHC diversity in a declining frog. Conserv Genet. 2018;19:309–322.

17. Cheng TL, Rovito SM, Wake DB, et al. Coincident mass extirpation of neotropical amphibians with the emergence of the infectious fungal pathogen *Batrachochytrium dendrobatidis*. Proc Natl Acad Sci U S A. 2011;108(23):9502–9507.

18. Azat C, Alvarado-Rybak M, Solano-Iguaran JJ, et al. Synthesis of *Batrachochytrium dendrobatidis* infection in South America: amphibian species under risk and areas to focus research and disease mitigation. Ecography. 2022;2022(7):e05977.

19. Coloma LA, Acosta-Buenaño NA, Acosta-Buenaño NA, et al. Anfibios de Ecuador [Amphibians of Ecuador]. Pichincha: Centro Jambatu; 2018.

20. Ortega-Andrade HM, Blanco MR, Cisneros-Heredia DF, et al. Red List assessment of amphibian species of Ecuador: A multidimensional approach for their conservation. PLoS One. 2021;16(5):e0251027.

21. Guayasamin JM, Mendoza AM, Longo AV, et al. High prevalence of *Batrachochytrium dendrobatidis* in an Andean frog community (Reserva Las Gralarias, Ecuador). Amphib Reptile Conserv. 2014;8:33–44.

22. Bresciano JC, Salvador CA, Paz-y-Miño C, et al. Variation in the presence of anti-*batrachochytrium dendrobatidis* bacteria of amphibians across life stages and elevations in ecuador. EcoHealth. 2015;12(2):310–319.

23. Menéndez-Guerrero PA, Graham CH. Evaluating multiple causes of amphibian declines of Ecuador using geographical quantitative analyses. Ecography. 2013;36(7):756–769.

24. Puschendorf R, Carnaval AC, VanDerWal J, et al. Distribution models for the amphibian chytrid *Batrachochytrium dendrobatidis* in Costa Rica: proposing climatic refuges as a conservation tool. Divers Distrib. 2009;15(3):401–408.

25. Becker CG, Rodriguez D, Lambertini C, et al. Historical dynamics of *Batrachochytrium dendrobatidis* in Amazonia. Ecography. 2016;39(10):954–960.

26. Deichmann JL, Bruce Williamson G, Lima AP, et al. A note on amphibian decline in a central Amazonian lowland forest. Biodivers Conserv. 2010;19(12):3619–3627.

27. McCracken S, Gaertner JP, Forstner MR, et al. Detection of *Batrachochytrium dendrobatidis* in amphibians from the forest floor to the upper canopy of an Ecuadorian Amazon lowland rainforest. Herpetol Rev. 2009;40(2):190.

28. Farrer RA, Weinert LA, Bielby J, et al. Multiple emergences of genetically diverse amphibian-infecting chytrids include a globalized hypervirulent recombinant lineage. Proc Natl Acad Sci U S A. 2011;108(46):18732–18736.

29. O’Hanlon SJ, Rieux A, Farrer RA, et al. Recent Asian origin of chytrid fungi causing global amphibian declines. Science. 2018;360(6389).

30. Mesquita AFC, Lambertini C, Lyra M, et al. Low resistance to chytridiomycosis in direct-developing amphibians. Sci Rep. 2017;7(1):16605.

31. Sette CM, Vredenburg VT, Zink AG. Differences in fungal disease dynamics in co-occurring terrestrial and aquatic amphibians. EcoHealth. 2020;17(3):302–314.

32. Altig R, McDiarmid RW. Morphological diversity and evolution of egg and clutch structure in amphibians. Herpetol Monogr. 2007;21(1):1–32.

33. Duellman WE, Trueb L. Biology of amphibians. Baltimore, Md.: JHU Press; 1994. en.

34. Bielby J, Cooper N, Cunningham Aa, et al. Predicting susceptibility to future declines in the world’s frogs. Conserv Lett. 2008;1(2):82–90.

35. Brem FMR, Lips KR. *Batrachochytrium dendrobatidis* infection patterns among Panamanian amphibian species, habitats and elevations during epizootic and enzootic stages. Dis Aquat Organ. 2008;81(3):189–202.

36. Ribeiro JW, Siqueira T, DiRenzo GV, et al. Assessing amphibian disease risk across tropical streams while accounting for imperfect pathogen detection. Oecologia. 2020;193(1):237–248.

37. Gründler MC, Toledo LF, Parra-Olea G, et al. Interaction between breeding habitat and elevation affects prevalence but not infection intensity of *Batrachochytrium dendrobatidis* in Brazilian anuran assemblages. Dis Aquat Organ. 2012;97(3):173–184.

38. Bass MS, Finer M, Jenkins CN, et al. Global conservation significance of Ecuador’s Yasuní National Park. PLoS One. 2010;5(1):e8767.

39. Tiputini U. 2023. Available from: https://fieldguides.fieldmuseum.org/sites/default/files/rapid-color-guides-pdfs/193_ecuador_ranas_de_la_amazonia_v3.pdf

40. Kriger KM, Hines HB, Hyatt AD, et al. Techniques for detecting chytridiomycosis in wild frogs: comparing histology with real-time Taqman PCR. Dis Aquat Organ. 2006;71(2):141–148.

41. Fisher MC, Stajich JE, Farrer R. Emergence of the chytrid fungus Batrachochytrium dendrobatidis and global amphibian declines. Evolution of virulence in eukaryotic microbes. Hoboken, NJ: Wiley; 2012. p. 461-472.

42. Rosenblum EB, James TY, Zamudio KR, et al. Complex history of the amphibian-killing chytrid fungus revealed with genome resequencing data. Proc Natl Acad Sci U S A. 2013 Jun 4;110(23):9385–9390.

43. Team’ RC. R: A language and environment for statistical computing Vienna, Austria: R Foundation for Statistical Computing; 2021. Available from: https://www.R-project.org/

44. Fox J, Weisberg S, Adler D, et al. Package ‘car’. Vienna: R Foundation for Statistical Computing. 2012;16.

45. Harrison E, Drake T, Ots R. Package “finalfit.”. Retrieved February. 2020;29:2020.

46. Lips KR. Mass mortality and population declines of anurans at an upland site in western panama. Conserv Biol. 1999;13(1):117–125.

47. Longcore JE, Pessier AP, Nichols DK. *Batrachochytrium dendrobatidis* gen. et sp. nov., a chytrid pathogenic to amphibians. Mycologia. 1999;91(2):219–227.

48. Ron SR. Predicting the distribution of the amphibian pathogen *batrachochytrium dendrobatidis* in the New World. Biotropica. 2005;37(2):209–221.

49. Liu X, Rohr JR, Li Y. Climate, vegetation, introduced hosts and trade shape a global wildlife pandemic. Proc R Soc B. 2013;280(1753):20122506.

50. Zumbado-Ulate H, García-Rodríguez A, Vredenburg VT, et al. Infection with *Batrachochytrium dendrobatidis* is common in tropical lowland habitats: Implications for amphibian conservation. Ecol Evol. 2019;9(8):4917–4930.

51. Russell ID, Larson JG, May Rv, et al. Widespread chytrid infection across frogs in the Peruvian Amazon suggests critical role for low elevation in pathogen spread and persistence. PLoS One. 2019;14(10):e0222718.

52. Rodríguez-Brenes S, Rodriguez D, Ibáñez R, et al. Spread of amphibian chytrid fungus across lowland populations of Túngara frogs in Panamá. PLoS One. 2016;11(5):e0155745.

53. von May R, Catenazzi A, Santa-Cruz R, et al. Microhabitat temperatures and prevalence of the pathogenic fungus *batrachochytrium dendrobatidis* in lowland amazonian frogs. Trop Conserv Sci. 2018;11:1–13.

54. Ruggeri J, Martins AGDS, Domingos AHR, et al. Seasonal prevalence of the amphibian chytrid in a tropical pond-dwelling tadpole species. Diseases of Aquatic Organisms. 2020;142:171–176.

55. Wilcoxen TE, Boughton RK, Schoech SJ. Selection on innate immunity and body condition in Florida scrub-jays throughout an epidemic. Biol Lett. 2010;6(4):552–554.

56. Lamirande EW, Nichols DK, editors. Effects of host age on susceptibility to cutaneous chytridiomycosis in blue-and-yellow poison dart frogs (*Dendrobates tinctorius*). Proceedings of the sixth international symposium on the pathology of reptiles and amphibians; 2001 Apr 18-19 St. Paul, MN.

57. Møller AP, Christe P, Erritzøe J, et al. Condition, disease and immune defence. Oikos. 1998;83(2):301–306.

58. Burrow AK, Rumschlag SL, Boone MD. Host size influences the effects of four isolates of an amphibian chytrid fungus. Ecol Evol. 2017;7(22):9196–9202.

59. Parris MJ, Cornelius TO. Fungal pathogen causes competitive and developmental stress in larval amphibian communities. Ecology. 2004;85(12):3385–3395.

60. Lips KR, Brem F, Brenes R, et al. Emerging infectious disease and the loss of biodiversity in a Neotropical amphibian community. Proc Natl Acad Sci U S A. 2006;103(9):3165–3170.

61. Sabagh LT, Ferreira RB, Rocha CFD. Host bromeliads and their associated frog species: Further considerations on the importance of species interactions for conservation. Symbiosis. 2017;73(3):201–211.

62. Tonini JFR, Ferreira RB, Pyron RA. Specialized breeding in plants affects diversification trajectories in Neotropical frogs. Evolution. 2020;74(8):1815–1825.

63. Peixoto OL. Associação de anuros a bromeliáceas na Mata Atlântica. RBCV. 2013;17(2):75–83.

64. Ruano-Fajardo G, Toledo L, Mott T. Jumping into a trap: high prevalence of chytrid fungus in the preferred microhabitats of a bromeliad-specialist frog. Diseases of Aquatic Organisms. 2016;121(3):223–232.

65. Burrowes PA, De la Riva I. Unraveling the historical prevalence of the invasive chytrid fungus in the Bolivian Andes: implications in recent amphibian declines. Biol Invasions. 2017;19(6):1781–1794.

66. Crawford AJ, Lips KR, Bermingham E. Epidemic disease decimates amphibian abundance, species diversity, and evolutionary history in the highlands of central Panama. Proc Natl Acad Sci U S A. 2010;107(31):13777–13782.

